# Selective estradiol sensitivity in 12Z human endometriotic epithelial cell line

**DOI:** 10.64898/2025.12.31.697205

**Authors:** Shohini Banerjee, Esha Chopra, Ian M. Smith, Kimberly M. Stroka

## Abstract

Estradiol (E2) is a potent estrogen molecule that plays a crucial role in regulating numerous healthy and pathophysiological processes. To model estrogen-dependent cellular activities *in vitro* in a physiologically meaningful way, cells chosen for experimentation should exhibit sensitivity to E2. The 12Z human endometriotic epithelial cell line is gaining attention as a model for estrogen-dependent conditions, but its functional responsiveness to E2 has not been well characterized. In the following study, we found that E2 does not influence 12Z proliferation, collective migration, or single-cell migration. However, 12Zs were selectively responsive to E2 in 3D migration models where the cells were physically confined. Upregulating ER⍰, the primary mediator of estrogenic action, in the 12Zs did not enhance their functional E2 sensitivity. To delineate the underpinning of these behaviors, RNA sequencing was performed and revealed a differential expression of pseudogenes and non-coding RNAs in E2-treated 12Zs compared to vehicle control-treated 12Zs. Several signal transduction genes were significantly downregulated in *ESR1*-overexpressing 12Zs compared to normal 12Zs, which may play a role in their persistent lack of E2 sensitivity. These results prompt us to question whether the 12Z cell line, which mostly lacks functional E2 responsiveness, should be used *in vitro* as a model of estrogen-dependent processes and conditions.

## INTRODUCTION

Estrogens are key hormone molecules that are intimately involved in numerous healthy and disease physiological systems, including but not limited to the reproductive system, cardiovascular health, immune activity, cancer metastasis, endometriosis, polycystic ovary syndrome, and uterine fibroids (1–4). Estradiol (E2), a potent estrogen, binds with estrogen receptors (ERs) located in the nucleus and on the plasma membrane to directly and indirectly regulate gene expression. This process impacts downstream cellular activities, such as cell proliferation, morphology, differentiation, migration, and invasion (5). When modeling estradiol-dependent biological processes *in vitro*, it is important to use cells that retain their hormone sensitivity. While the advent of cell lines is advantageous for widespread use in high-throughput experiments, their physiological relevance can be limited by the loss of primary cell traits. The 12Z human endometriotic epithelial cell line has been developed as a model of endometriosis and estrogen-dependent metastasis and shares some molecular similarities with primary cells (6,7). Although the cell line is gaining popularity, its responsiveness to E2 has not been well characterized. Given that endometriosis is marked by excessively high local levels of E2 that are known to impact primary cell behaviors (8–13), the 12Z cell line should retain a similar sensitivity to E2. 12Z cells have been previously reported to express nuclear ERs (6,7), so we hypothesized that they would exhibit E2 dependence *in vitro*.

In the following study, we tested and characterized the E2 sensitivity of the 12Z cell line in a variety of proliferation and migration models. We found 12Zs exhibit selective responsiveness to E2, with altered behaviors in confined migration models but no changes seen in unconfined proliferation and migration systems. Moreover, increasing levels of the gene encoding ER⍰, a major mediator of estrogenic action (14), did not enhance 12Z responsiveness to E2. We used RNA sequencing to further inform these differences in E2 sensitivity. Given the insufficient responsiveness to E2, we question whether the 12Z cell line is appropriate for use in studying estrogen-dependent processes *in vitro*.

## MATERIALS AND METHODS

### Cell culture and treatments

The 12Z cell line was purchased from Applied Biological Materials (Cat. no. T0764). Cells were cultured in DMEM/F-12 (Gibco) supplemented with 10% fetal bovine serum (FBS) (Gibco) and 1% penicillin-streptomycin (Gibco). About 24-48 hours prior to experiments, the media was replaced with phenol-free DMEM/F-12 (Gibco) supplemented with 10% charcoal-stripped FBS (Gibco) and 1% penicillin-streptomycin (Gibco). A modified 12Z cell line with an upregulation of *ESR1* (*ESR1*-12Z) was generously gifted by Dr. Fazleabas’s lab and cultured in phenol-free DMEM/F-12 supplemented with 10% charcoal-stripped FBS and 1% penicillin-streptomycin. 17β-estradiol (E2) was purchased from Sigma Aldrich (Cat. # E8875-1G) and dissolved in 100% ethanol to create a stock solution. Another vial of E2 was purchased from MP Biomedicals (Cat. no. 02194565-CF) and dissolved in 100% ethanol to create a stock solution (denoted as MPB E2). Cells were treated with media containing a vehicle (0.001% ethanol), 10 nM E2, or 10 nM MPB E2, as well as other treatments as specified in the Results section for the dose-dependent experiments. Unless specified as MPB E2, E2 refers to the batch sourced from Sigma Aldrich.

### Proliferation and wound healing assays

To quantify cellular proliferation, a Cell Counting Kit-8 assay was employed. Briefly, 10,000 cells were seeded per well in a 96-well plate and cultured for 48 hr with treatments. These include the mitotic inhibitor, mitomycin C (MMC, Sigma Aldrich, cat. #M5353) which was used as a positive control at a concentration of 10 µg/mL. 10 µL of CCK-8 solution was then added to each well and incubated for 1-1.5 hr at 37°C. Absorbance measurements were taken in a microplate reader at 450 nm. Plotted absorbance values were normalized to the vehicle group as specified in each figure. To measure collective cell migration, scratch wound healing assays were performed. Approximately 90,000 cells were seeded per well in a 24-well plate coated with 10 µg/mL collagen I from rat tail (Corning, cat. # 354249). Cells were incubated for 48 hr with treatments as they grew into confluent monolayers. Monolayers were manually scratched with a p-200 pipette tip and rinsed with phosphate-buffered saline (PBS) before replacing media with treatments. Wounds were imaged under phase contrast at 10X one hour post-scratch (Initial wound area) and 7.25 hr post-scratch (Final wound area). Wound closure was calculated as (Initial wound area – Final wound area) / Initial wound area, with wound areas determined using ImageJ (National Institutes of Health, Bethesda, MD, USA.

### Quantification of cellular morphology

Morphological parameters including cell area, perimeter, solidity, and roundness were extracted using ImageJ to manually trace cells from phase contrast microscope images. Solidity was defined as 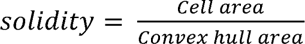 where a cell that is perfectly circular would have a value of 1.0 and a cell that has protrusions and/or indentations would have a value closer to 0. Roundness was defined as 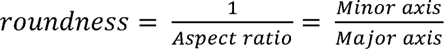, where a highly elongated cell would have a value close to 0.

### Single cell random migration analysis

The random migration of single cells was analyzed from time-lapse phase contrast images. Briefly, approximately 5,000 cells were seeded per well in a 24-well plate coated with 10 µg/mL rat tail collagen I, except for the positive control, where the wells were coated with 20 µg/mL poly-D lysine (PDL) to prevent integrin-based cell attachment. Cells were incubated with treatments for 48 hr and then imaged under phase contrast at 10X for 12 hr, with a time step of 10 min. Individual cell trajectories were tracked using the Manual Tracking plugin in ImageJ by clicking on the estimated geometric center of the cell at each time point. Subsequent analysis was performed in MATLAB. Average speed was defined as the cell’s total path length divided by the duration of the time-lapse. Data represent speed normalized to the vehicle. Directional persistence was defined as the cell’s end-to-end displacement divided by its total path length, where a cell that progresses in a perfectly straight line from point A to point B would have a directional persistence of 1; a cell that originates at a point A, moves in a straight line to point B, and directly returns to point A would have a directional persistence of 0. Mean-square displacement (MSD), which is a proxy for how much area (in µm^2^) a cell covered during its trajectory, is defined as MSD == (r^2^) == 4DΔt, where Δt is the number of time steps (Δt == 1 represents one step, or 10 min; Δt == 2 represents two time steps, or 20 min, etc.) and D represents the diffusivity (µm^2^/time step). Diffusivity was derived from the slope m of the linear portion of the MSD curve, where m == 4D and 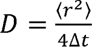.

### Transwell migration assay

Cell transmigration through a semi-permeable membrane was performed using a transwell migration assay. Transwell inserts for a 24-well plate were coated with 10 µg/mL collagen I from rat tail. 100,000 cells were seeded in the upper chamber in serum-free media, with the lower chamber containing media supplemented with 10% FBS. Cells were allowed to transmigrate for 24 hr prior to quantification. Then, media was aspirated from the upper chambers, the top of the transwell inserts were swabbed gently, and the inserts were stained with 5 µM calcein AM for 30 min. Transmigrated cells were imaged under fluorescence. Data are normalized to the vehicle.

### Cell migration through confined microfluidic channels and analysis

To analyze 12Z migration in narrow spaces, cells were sent through microfluidic devices containing narrow channels (schematic in Fig. 5B). As previously defined (15), microfluidic devices with confining microchannels 3 µm wide x 10 µm tall x 200 µm long were cast in polydimethylsiloxane (PDMS, Dow) using a positive mold created by photolithography. The inlets and outlets of the PDMS device were made with a 4 mm biopsy punch. The devices were then irreversibly adhered to a #1 glass coverslip via plasma treatment, followed by 5 min of pressure and an overnight bake at 80°C. One hour before their use, the devices were again plasma treated to create hydrophilic surfaces, and 0.1 mg/mL type I collagen (cat. # C3867, Sigma Aldrich) in PBS was added to the inlets to functionalize the inner walls. Devices with collagen were incubated for 1 hour at room temperature. The devices were then added to a 4-well plate (07-000-207, Fisher Scientific) and UV-treated for 10 min to sterilize them before adding cells. Following initial cell splitting, cells were resuspended in serum-free media, counted, and centrifuged again. Cells were then resuspended to a concentration of 100,000 cells/20 µL of serum-free media containing vehicle or E2 treatments. 20 µL of this cell suspension was added to the microchannel cell inlet. After adding cells, the devices were placed in an incubator for 10 min to ensure cell adhesion to the microchannel walls. Afterwards, excess media and adhered cells were removed, and media was added to the remaining wells. Cell inlets and outlets, in addition to 2 media inlets and the media outlet, were filled with serum-free media. The final media inlet was filled with full media to induce a chemokine gradient. Approximately ten regions of interest were selected for each device and imaged at 20X under phase contrast every 15 min for 16 hours.

Individual cell trajectories while in the microchannels were tracked using the Manual Tracking PlugIn in ImageJ by clicking on the nucleus of the cell at each time point. Subsequent analysis was performed in MATLAB. Speed was defined as the cell’s total path length divided by the duration of the time-lapse. Data represent speed normalized to the vehicle. Directional persistence was defined as the cell’s end-to-end displacement divided by its total path length, where a cell that progresses in a perfectly straight line from point A to point B would have a directional persistence of 1; a cell that originates at a point A, moves in a straight line to point B, and directly returns to point A would have a directional persistence of 0.

### Spheroid outgrowth assay

A spheroid outgrowth experiment was performed to quantify the expansion of 3D cell spheroids on a collagen I gel. 4,000 12Z cells were seeded per well in a round-bottom 96-well ultra-low attachment plate and centrifuged at 150xg for 2 min. Spheroids were harvested at 48 hr and seeded on top of collagen I gels. Phase contrast images were taken at time of seeding (Day 0) and 24 hr (Day 1). The projected area of each spheroid was measured in ImageJ; data represent the area on Day 1 relative to the initial area on Day 0.

### Western blotting

Relative levels of estrogen receptor-α (ERα) were measured with western blotting to validate the overexpression of its encoding gene, *ESR1*, in the *ESR1*-12Zs. Cells were cultured for 48 hr and lysed with a protease cocktail inhibitor (1:100; Sigma Aldrich) and phenylmethylsulphonyl fluoride (1:500; Sigma Aldrich, cat. #P7626) in Pierce RIPA buffer (ThermoFisher Scientific). A BCA Protein Assay Kit (ThermoFisher Scientific) was used to determine protein concentration in the lysates. Samples were mixed 1:1 with a buffer containing 20:1 2X Laemmli to β-mercaptoethanol (Bio-Rad, cat. #1610737 and #1610710, respectively) and boiled. Equal amounts of protein were loaded into a Mini-PROTEAN TGX 4-20% precast gel (Bio-Rad) and separated by sodium dodecyl sulfate-polyacrylamide gel electrophoresis. Samples were then transferred to a PVDF membrane via the Trans-Blot Turbo machine (Bio-Rad) on the Mixed MW (Turbo) protocol setting (1.3A, up to 25V, 7 min). PVDF membranes were blocked in an Intercept (PBS) blocking buffer (LI-COR) for one hour and incubated overnight at 4°C with a primary antibody solution consisting of anti-ERα (1:500; Santa Cruz Biotechnology) and anti-α-smooth muscle actin (1:500; Sigma Aldrich, cat. #A5228) as a control in a 1:1 Intercept blocking buffer to 0.1% Tween 20 in PBS mix (PBST). The next day, membranes were washed with PBST and incubated for one hour at RT with a secondary antibody solution consisting of IRDye 680RD Goat anti-Rabbit and Goat anti-Mouse (each at 1:10,000; LI-COR, cat. #926-68071 and #926-68070, respectively) in a 1:1 Intercept blocking buffer to PBST mix. Membranes were then washed with PBS, visualized on an Odyssey CLx imager, and analyzed with the Image Studio software (LI-COR).

### RNA-sequencing and data analysis

Vehicle- and E2-treated 12Zs as well as non-treated 12Zs and non-treated *ESR1*-12Z samples were sent for RNA extraction, integrity analysis, library preparation, and sequencing (Novogene Co, Ltd.). Three independent samples for each cell group were pooled before shipping. Messenger RNA was purified from total RNA using poly-T oligo-attached magnetic beads. After fragmentation, the first strand cDNA was synthesized using random hexamer primers. Then the second strand cDNA was synthesized using dUTP, instead of dTTP. The directional library was ready after end repair, A-tailing, adapter ligation, size selection, USER enzyme digestion, amplification, and purification. The library was checked with Qubit and real-time PCR for quantification and bioanalyzer for size distribution detection. After library quality control, libraries were subjected to Illumina Novoseq 6000 sequencing (paired-end 150 bp) with data output of over 20 million read pairs per sample. Raw reads (fastq format) were processed through fastp software. Clean reads were obtained by removing reads containing adapter, reads containing ploy-N and low-quality reads from raw data. Full RNA sequencing methods are provided in the Supplementary Material (S1).

### Microscopy

For all phase contrast images and transmigration fluorescence images, live cells were imaged using the 10X objective (unless otherwise specified) on an Olympus IX-83 inverted microscope (Olympus, Center Valley, PA, USA) and the Olympus cellSens software. Cells were maintained at 37°C, 65% humidity, and 5% CO_2_ in an enclosed chamber surrounding the microscope.

### Statistical analysis

GraphPad Prism 10 (La Jolla, CA, USA) was utilized to generate graphs and perform statistical analyses. Outliers were identified using the ROUT method with a Q (maximum desired false discovery rate) = 1% only to remove definite outliers. Data were tested for normality. For comparisons between vehicle and E2 groups, unpaired Student’s *t*-tests were performed. For comparisons between three or more groups, ordinary one-way ANOVA with a Tukey’s multiple comparisons test was performed. Corresponding non-parametric tests were used for non-normally distributed datasets. For all tests, P values < 0.05 were considered to be statistically significant. Data are represented as mean ± standard error of the mean. All experiments were performed in at least triplicates.

## RESULTS

### Determination of E2 incubation time and concentration for 12Zs

Multiple functional assays were employed to determine the optimal incubation time and concentration of E2 to apply to 12Z cells. To determine the most effective dose of E2, cells were treated with 10^−9^ M, 10^−8^ M, 10^−7^ M, or 10^−6^ M E2 for 48 hr before testing their migratory and proliferative response. Instead of including all the corresponding vehicle controls, only the lower and upper extremes were tested (0.0001% EtOH corresponding to 10^−9^ M E2 and 0.1% EtOH corresponding to 10^−6^ M E2). None of the E2 concentrations impacted wound closure compared to either of the vehicle controls (Fig. 1A). In a CCK-8 proliferation assay, 10^−9^ M and 10^−8^ M E2 treatments each led to a non-significant increase in absorbance compared to the 0.0001% and 0.1% EtOH control groups (Fig. 1B). This result, combined with support from the literature (9–11,13,16,17), led us to select 10^−8^ M (10 nM) as our concentration for all subsequent experiments involving E2. To determine the most effective incubation time with chosen treatments (0.001% EtOH vehicle control and 10 nM E2) for the 12Z cells, we analyzed cell morphology at the time of adding treatments (t = 0), 24 hr, 48 hr, and 72 hr from phase contrast images. Results show that E2 induces slight changes in cell morphology after 48-72 hr (Fig. 1C-G). Cell area (Fig. 1C) and perimeter (Fig. 1D) decreased at 72 and 48 hr, respectively. There were no changes in solidity across the incubation times (Fig. 1E) but a decrease in roundness at 48 hr (Fig. 1F). Representative images of 12Z cells treated for 48 hr are shown (Fig. 1G). Given the decrease in cell perimeter and roundness at 48 hr, along with support from literature (10,13,17,18), we chose 48 hr as the incubation time for all subsequent experiments involving E2.

**Figure 1.**
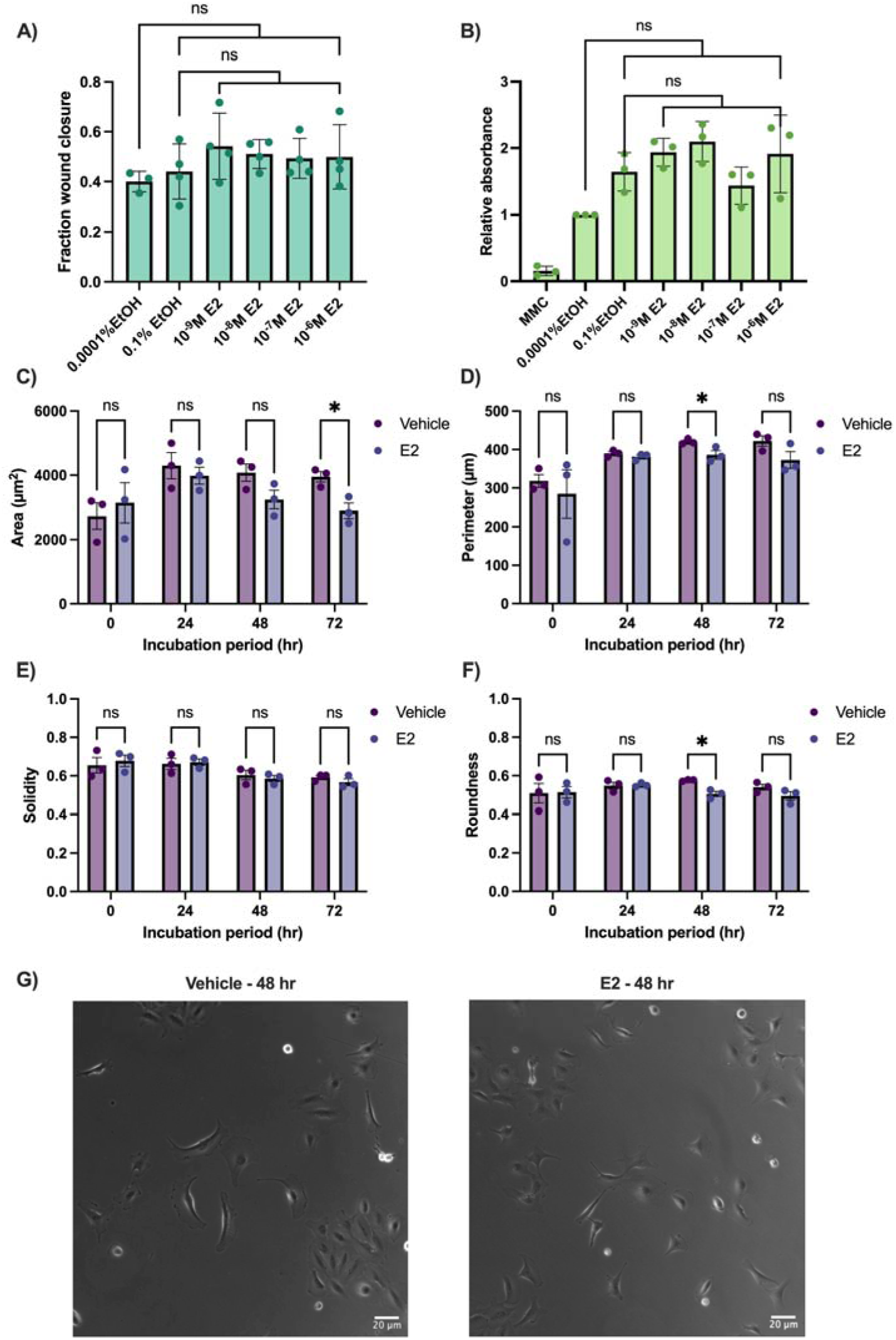
12Z characterization and determination of E2 dose and incubation time. **A)** Cell migration in a wound healing assay and **B)** absorbance from a CCK-8 assay, reported relative to the the 0.0001% EtOH group (with mitomycin C as a positive control), were quantified. From this point on, all subsequent experiments were performed using a concentration of 10 nM E2 and a corresponding 0.001% EtOH vehicle control. We then imaged cells treated with this vehicle control and 10 nM E2 for multiple incubation times and quantified cell morphology parameters including **C)** area, **D)** perimeter, **E)** solidity, and **F)** roundness. **G)** Representative images from the 48hr incubation period are shown. From this point on, 48 hr was selected as the treatment incubation period for all subsequent experiments. Scalebars = 20 µm. Data points represent biological replicates, error bars represent standard error of the mean. *P ≤ 0.05, **P ≤ 0.01. EtOH; ethanol. E2; estradiol. MMC; mitomycin C.

### E2 does not affect 12Z proliferation, collective migration, or single-cell random migration

The responsiveness of 12Zs to 10 nM E2 treatment for 48 hr was assessed *in vitro* with proliferation and migration assays. E2 had no influence on 12Z proliferation in a CCK-8 assay, in which MMC was used as a positive control and absorbance values were plotted relative to the vehicle control group (Fig. 2A). E2 also had no impact on collective migration in a wound closure assay (Fig. 2B,C). We then assessed single-cell random migration in response to E2 (Fig. 2D-H) and used a poly-D lysine (PDL) coating to prevent integrin-based cell-substrate attachment as a positive control. Individual cell trajectories show that E2-treated cells covered slightly more area than vehicle treated cells (Fig. 2D). E2 did not impact cell speed (Fig. 2E), directional persistence (Fig. 2F), or diffusivity (Fig. 2G) compared to the vehicle control group. The mean-square displacement (MSD) graph shows that the E2-treated cells traversed the largest area, but not significantly more than the vehicle-treated cells (Fig. 2H). The 12Z cell line did not appear to be sensitive to E2 treatment in proliferation and 2D migration assays *in vitro*. In case our E2 sourced from Sigma Aldrich was defective, we decided to additionally test an alternative source of E2 purchased from MP Biomedicals and denoted as MPB E2. However, MPB E2 had no effect on 12Z wound closure migration (Supplementary Material Fig. S2A).

**Figure 2.**
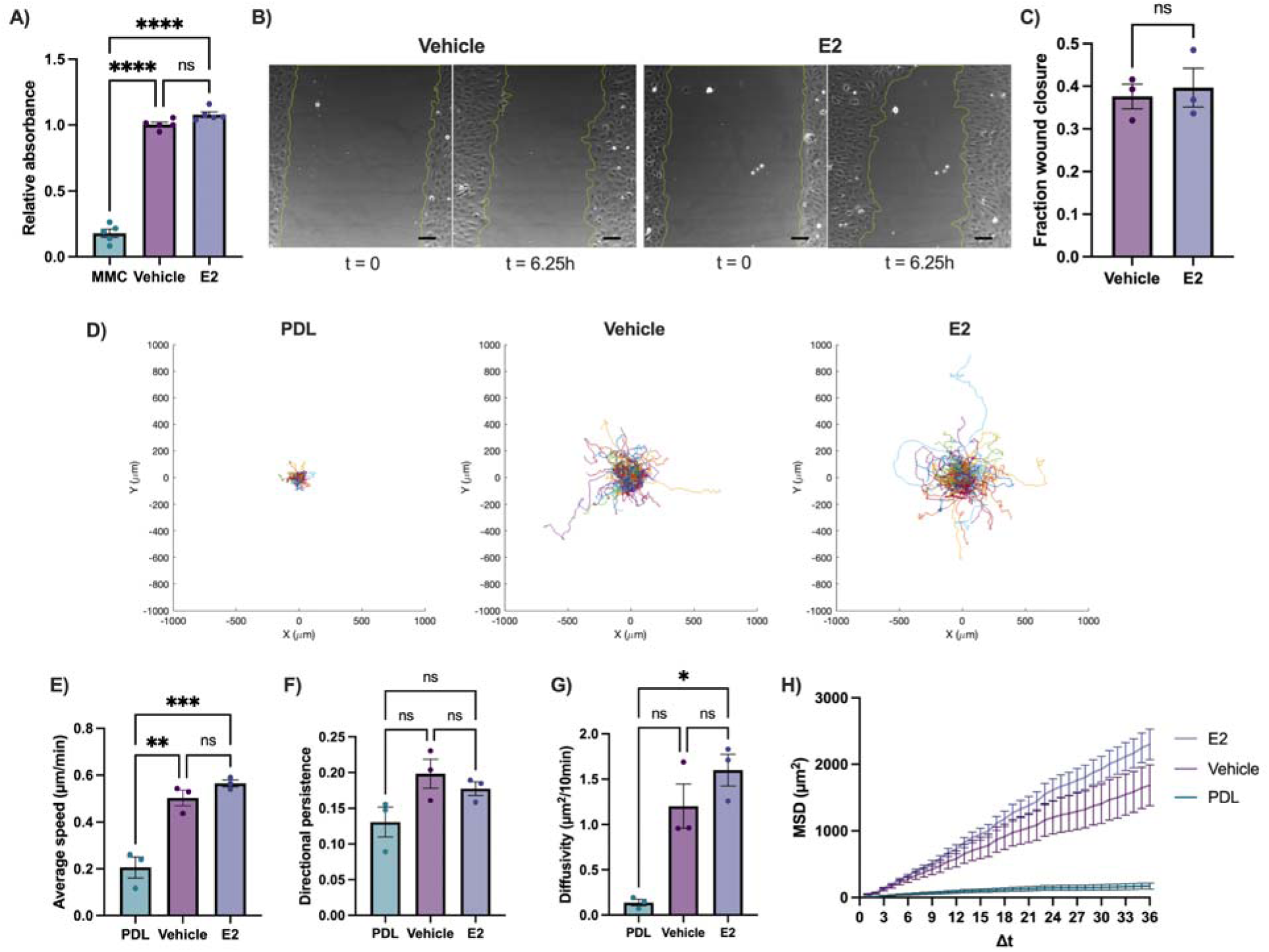
Effect of E2 on proliferative and migratory behaviors of 12Zs. 12Z cells were treated with 10 nM E2 for 48 hr and their functional behaviors were assessed. **A)** Proliferation was measured with a CCK-8 assay (with 10 µg/mL MMC as a positive control), and absorbance values are plotted relative to the vehicle control group. **B)** Representative pictures of wound closure are shown from time of scratch at t = 0 and midway through the experiment at t = 6.25 hr for both vehicle and E2-treated 12Zs. **C)** Fraction wound closure is calculated. **D–H)** Random migration was measured via single-cell tracking and a poly-D lysine coating, which prevents integrin-based attachment, was used as the positive control. **D)** Single-cell trajectories are shown. **E)** Speed, **F)** directional persistence (defined as the cell’s total displacement divided by the cell’s total path length), **G)** diffusivity, and **H)** MSD over increasing time steps are plotted. Scalebars = 100 µm. Data points represent biological replicates, error bars represent standard error of the mean. *P ≤ 0.05, **P ≤ 0.01, ***P ≤ 0.001, ****P ≤ 0.0001. MMC; mitomycin C. E2; estradiol. PDL; poly-D lysine.

### E2 differentially affects 12Z migration in confined and unconfined systems

The 12Z cell line showed a lack of sensitivity to E2 in proliferation and 2D migration assays, but cells are known to behave in a context-dependent manner: this could include differences between confined vs unconfined systems or between 2D vs 3D (19–22). After treating 12Zs with 10 nM E2 for 48 hr, their migration in confined and unconfined systems was analyzed (Fig. 3). E2 significantly increased transmigration in a transwell assay, with representative images showing the bottom-facing surfaces of the 8 µm-pore transwell inserts (Fig. 3A). Transmigration was quantified relative to the vehicle control (Fig. 3B). The effect of E2 was consistent when we tested the alternative source of E2, MPB E2, on 12Z transmigration (Supplementary Material Fig. S2B). In another confined migration model, we seeded 12Z cells in a previously defined PDMS microfluidic device containing narrow, 3-µm wide by 10-µm tall microchannels (Fig. 3C) (15). E2 increased cell migration speed through the channels (Fig. 3D) but did not impact chemotactic index, defined as the end-to-end displacement divided by the total cell path length (Fig. 3E). To model unconfined 3D migration, we seeded 12Z spheroids onto collagen I gels (Day 0) and measured their outgrowth 24 hr later (Day 1) (Fig. 3F). E2-treated spheroids did not exhibit differences in outgrowth compared to vehicle-treated spheroids (Fig. 3G).

**Figure 3.**
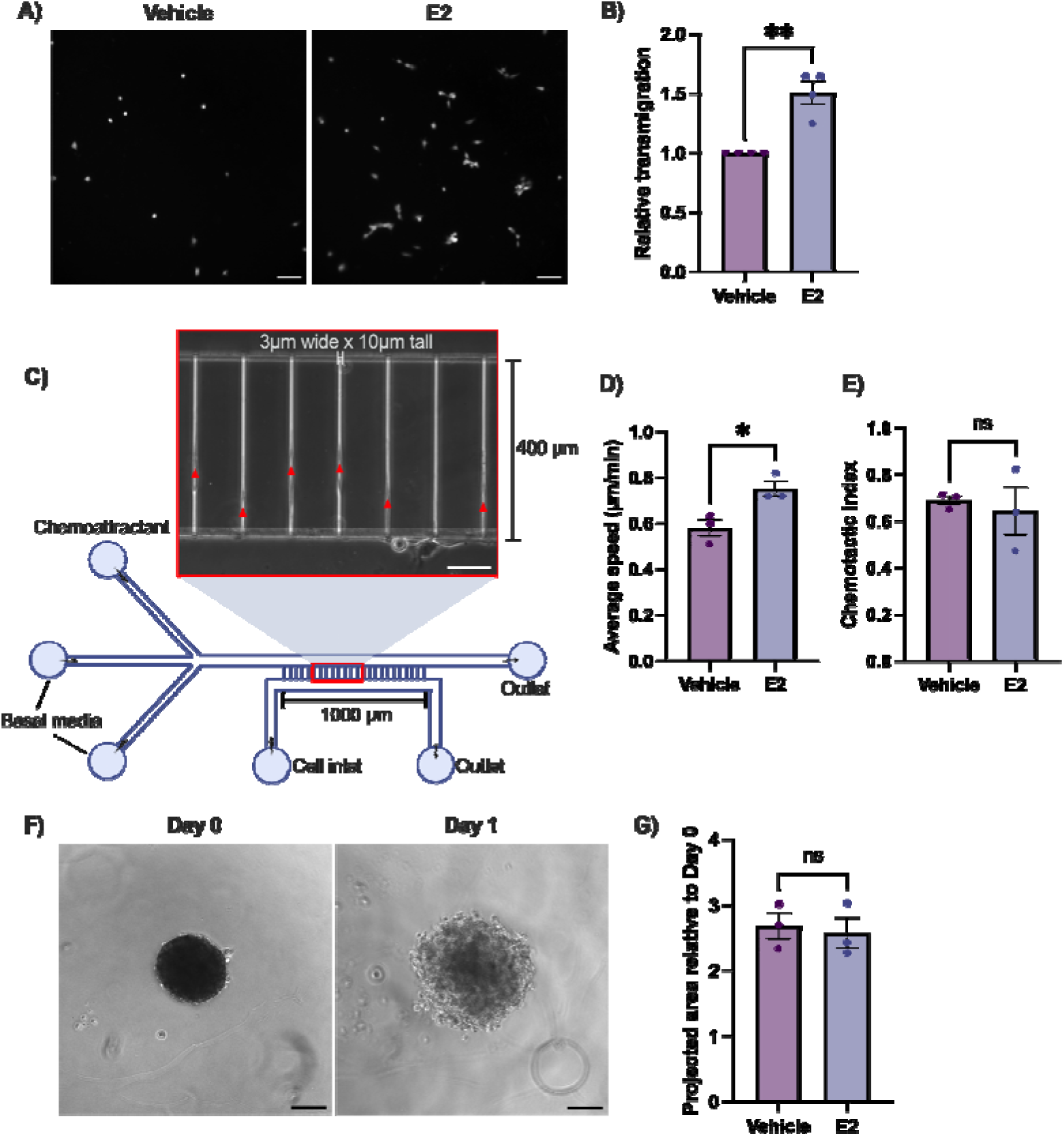
Impact of E2 on 12Zs in confined and unconfined systems. 12Z cells were treated with the vehicle or E2 for 48 hr prior to experiments. **A)** Transmigration in a transwell assay was measured and **B)** quantified relative to the vehicle control. **C-E)** Spontaneous migration of 12Zs through confining microchannels was observed. **C)** The design of the microfluidic device is depicted in the schematic which contains a representative image of cells moving through the microchannels towards the chemoattractant (media with 10% charcoal-stripped FBS); the cell nuclei are marked by red triangles. **D)** Cell speed and **E)** chemotactic index, defined as the end-to-end displacement divided by the total cell path length, were quantified. **F)** 3D 12Z spheroids were grown and seeded on collagen I gels (Day 0) and their projected area of outgrowth was measured the next day (Day 1). **G)** Projected area is calculated relative to Day 0. Scalebars = 100 µm in panels A and F and 50 µm in panel C. Data points represent biological replicates, error bars represent standard error of the mean. *P ≤ 0.05, **P ≤ 0.01. E2; estradiol.

### Upregulation of *ESR1* does not induce functional E2 sensitivity in the 12Zs

Our results show that the 12Z cell line is unresponsive to E2 treatment in most standard *in vitro* models of proliferation and unconfined migration. Estrogenic action is thought to be predominantly mediated by *ESR1,* the gene that encodes ER⍰ (14). We hypothesized that increasing the levels of ER⍰ would enhance the responsiveness of 12Zs to E2. We obtained a 12Z cell line genetically modified to upregulate *ESR1* (23) from Dr. Fazleabas’s lab (Michigan State University), hereafter denoted as *ESR1*-12Z. Upon receiving the *ESR1*-12Z cell line, we verified the increased expression of ER⍰ with a western blot (Fig. 4A). Western blot replicates and the effect of E2 on ER⍰ expression are reported (Supplementary Material S3). The *ESR1*-12Zs significantly overexpress ER⍰ by about 26-fold compared to the normal 12Zs; in fact, ER⍰ appears to be nearly absent in the 12Zs (Fig. 4B). However, after treating the *ESR1*-12Zs with 10 nM E2 (an alternate source of E2, denoted MPB E2, was also tested) for 48 hr, their functional behaviors showed no changes (Fig. 4C-E) compared to the vehicle control. In a CCK-8 proliferation assay with MMC as a positive control, neither source of E2 led to a change in absorbance compared to the vehicle control (Fig. 4C). Similarly, neither source of E2 significantly impacted wound closure migration (Fig. 4D) or migration through a transwell insert (Fig. 4E).

**Figure 4.**
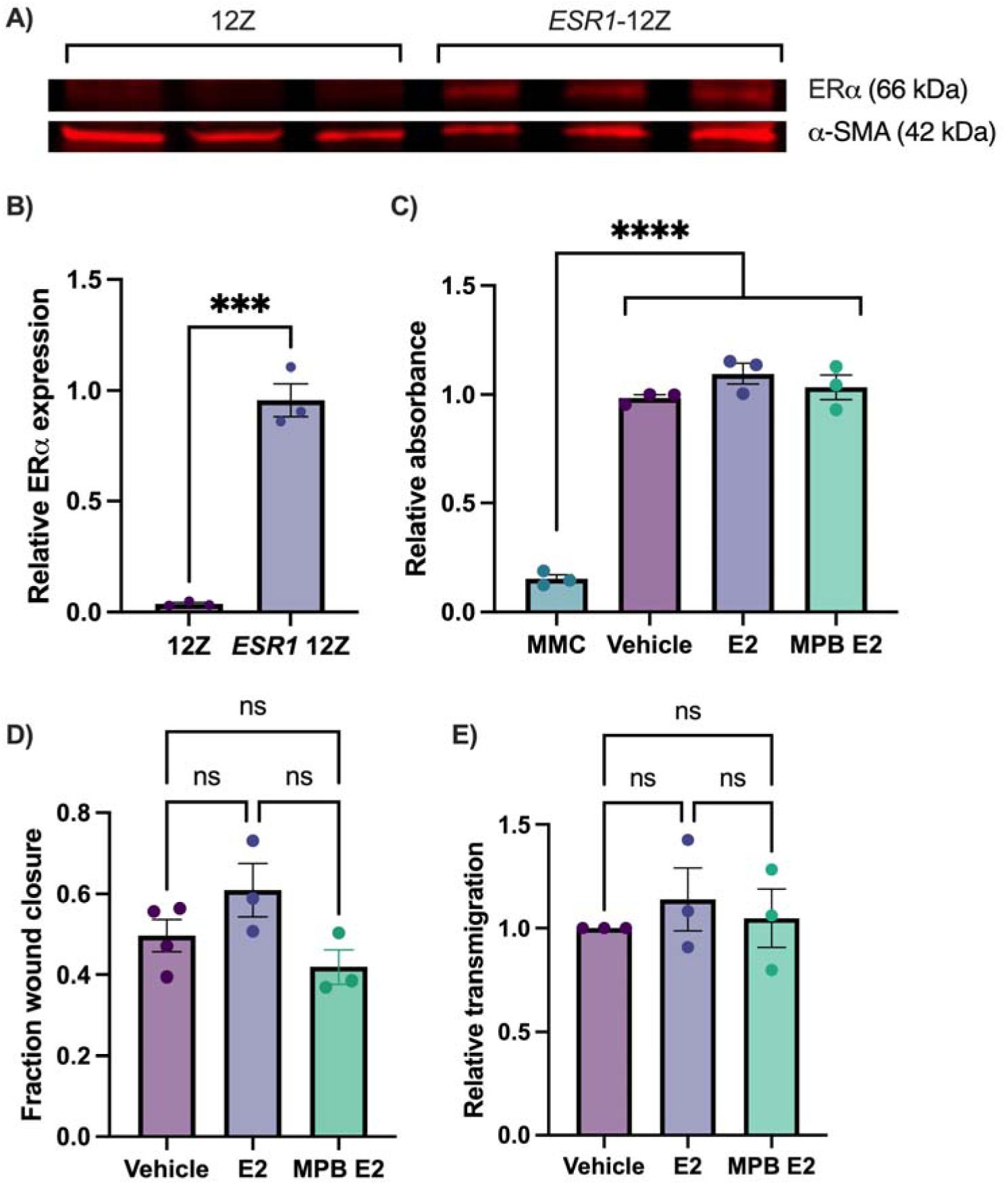
Effect of E2 on proliferative and migratory behaviors of *ESR1*-12Zs. We acquired 12Z cells stably transfected with *ESR1* (*ESR1*-12Zs) (23), the gene encoding the estrogen receptor ⍰ (ER⍰). **A)** A western blot for ER⍰ expression was performed using ⍰-smooth muscle actin (⍰-SMA) as the reference protein and **B)** quantified relative to ⍰-SMA to verify the upregulation of *ESR1* in the *ESR1*-12Zs. **C-E)** Then, *ESR1*-12Zs were subject to vehicle, 10 nM E2, or 10 nM MPB E2 for 48 hr and tested for **C)** proliferation in a CCK-8 assay (with 10 µg/mL MMC as a positive control) reported relative to the vehicle control, **D)** wound closure, and **E)** transwell migration reported relative to the vehicle control. Data points represent biological replicates, error bars represent standard error of the mean. ***P ≤ 0.001, ****P ≤ 0.0001. E2; estradiol (original batch purchased from Sigma Aldrich). MPB E2; MP Biomedicals estradiol. MMC; mitomycin C.

### Gene expression profiles: effects of E2 treatment or ESR1 upregulation

Vehicle-treated 12Zs (VE_12Z), E2-treated 12Zs (E2_12Z), normal 12Zs (NT_12Z), and *ESR1*-12Zs (ESR1_12Z) were sent for RNA extraction and sequencing (Novogene Co, Ltd.) to inform trends seen in E2 sensitivity in functional assays. Mainstream hierarchical clustering analysis shows some transcriptomic similarity between VE_12Z and E2_12Z (11,512 genes co-expressed, 451 and 580 genes uniquely expressed, respectively) and relatively less similarity between NT_12Z and ESR1-12Z (10,818 genes co-expressed, 1034 and 1065 genes uniquely expressed, respectively) (Fig. 5A). These intergroup differences are also shown in the 2D principal component analysis (Fig. 5B).

**Figure 5.**
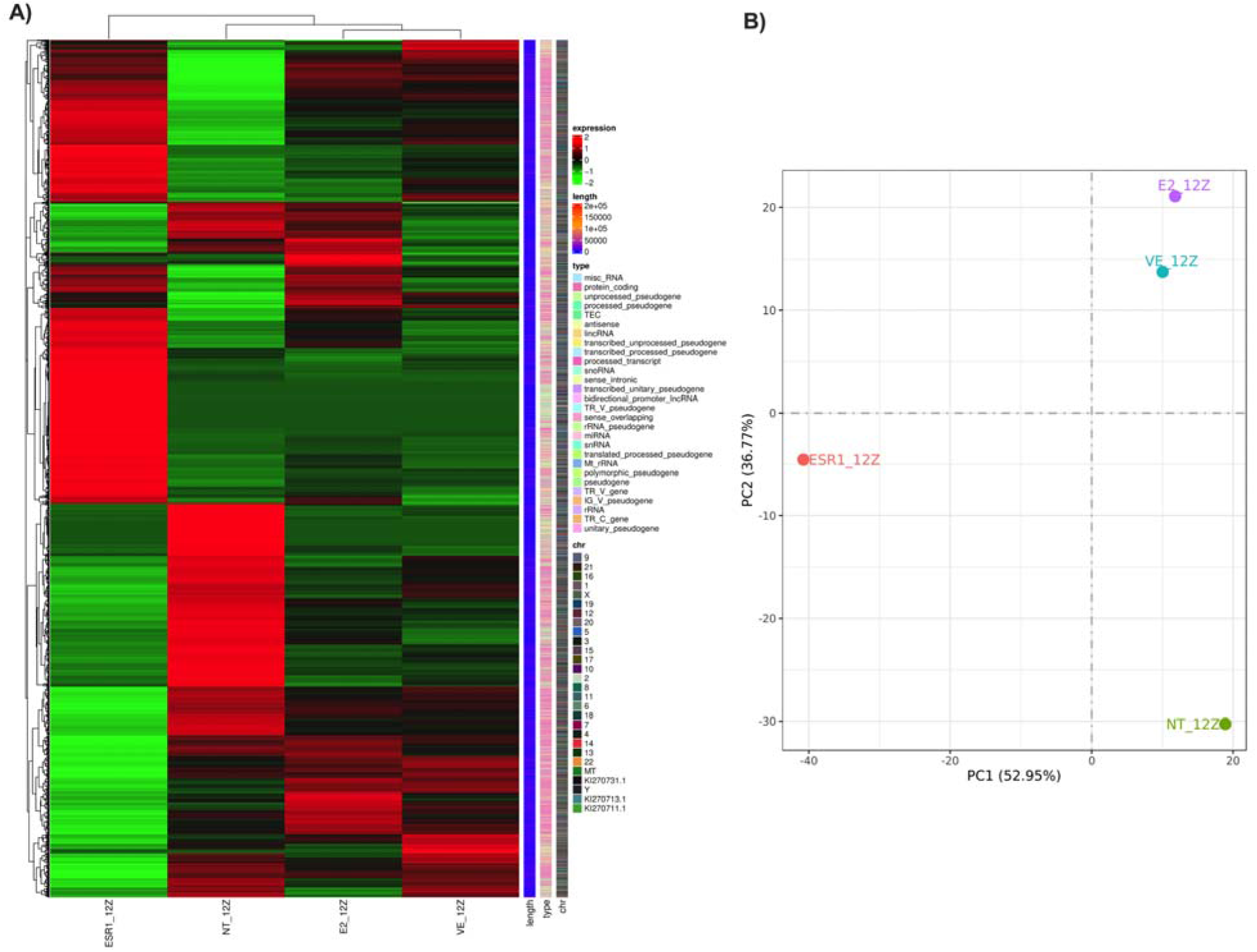
Gene expression profiles of vehicle-treated 12Zs, E2-treated 12Zs, normal 12Zs, and *ESR1*-12Zs. Vehicle-treated 12Zs (VE_12Z), E2-treated 12Zs (E2_12Z), normal 12Zs (NT_12Z), and *ESR1*-12Zs (ESR1_12Z) were sent for RNA extraction and sequencing (Novogene Co, Ltd.). **A)** Gene expression profiles can be visualized in the heatmap of mainstream hierarchical clustering and **B)** intergroup differences can be seen in the 2D principal component analysis plot. Each cell sample contains 3 trials pooled prior to sequencing.

A differential gene expression analysis was carried out using the EdgeR trimmed mean of M values method (24) (adjusted P < 0.05) to normalize the data. P-values were adjusted using the Benjamini-Hochberg method to control the error discovery rate (25). In the E2_12Z group compared to the VE_12Z group, there were 289 differentially expressed genes (DEGs): 185 upregulated and 104 downregulated (Supplementary Material S4). We found that E2 induced a differential expression of many pseudogenes and other non-coding RNAs, as annotated on the volcano plot (Fig. 6A). Out of the top 10 DEGs, 7 were pseudogenes or non-coding RNA. *RN7SL5P*, *PKD1P1*, and *AL591846.1* are examples of significantly upregulated pseudogenes in E2_12Z compared to VE_12Z, while non-coding RNAs *AC126755.2*, *AL391280.1*, *AC008555.8*, and *AC010331.1* were significantly downregulated (Fig. 6B). The roles of these pseudogenes and non-coding RNAs are not well defined, although they are likely capable of indirectly regulating gene expression (26,27).

**Figure 6.**
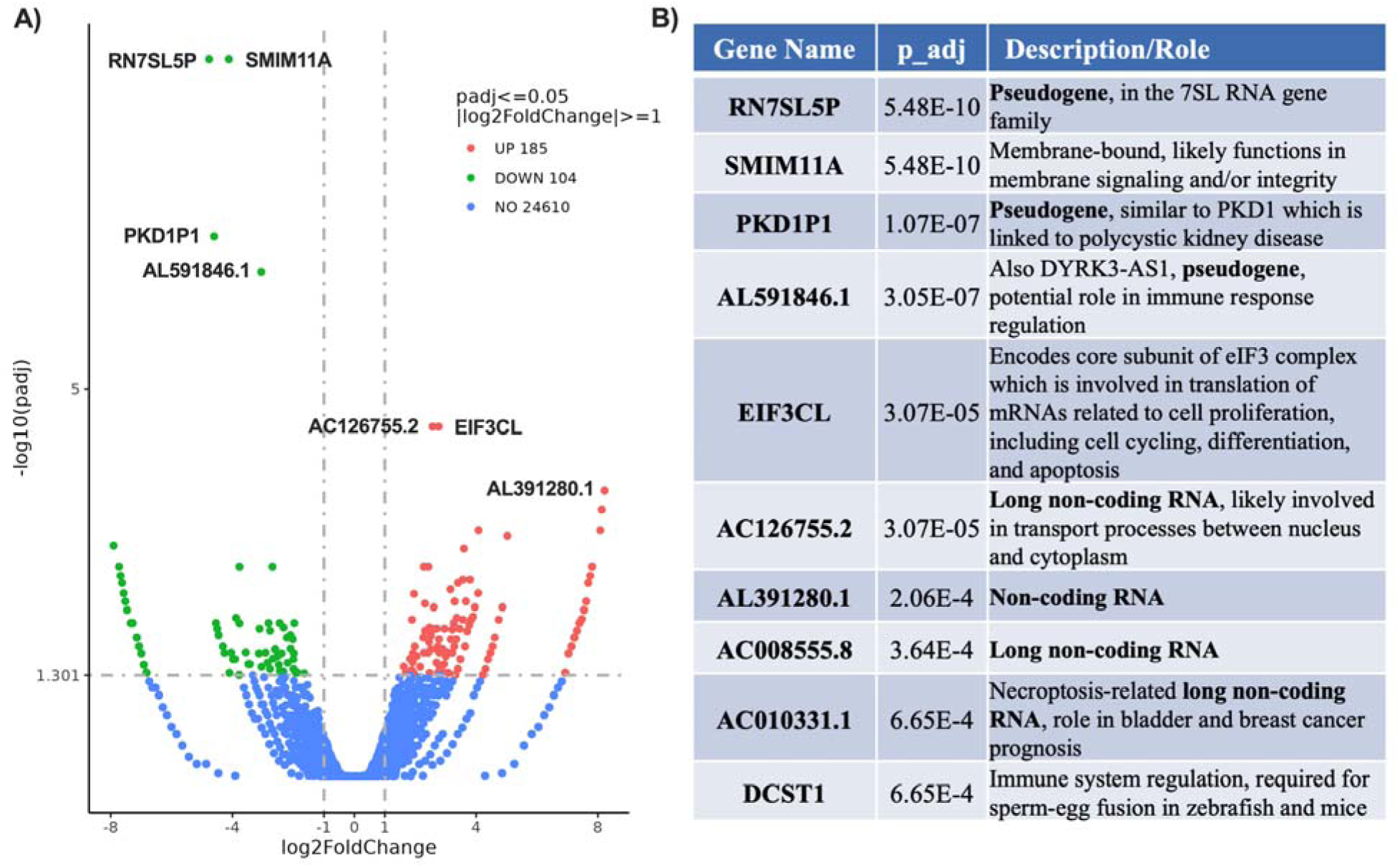
Differentially expressed genes between 12Zs treated with vehicle or E2. **A)** Several top DEGs are annotated on the volcano plot. **B)** The top 10 significantly DEGs, including several pseudogenes and non-coding RNAs, are tabulated and described along with their adjusted p-values (p_adj). Differential gene expression analysis was carried out using EdgeR’s trimmed mean of M values with p_adj < 0.05 (24). P-values were adjusted using the Benjamini-Hochberg method to control the error discovery rate (25).

The differential gene expression analysis between NT_12Z and ESR1_12Z revealed much larger transcriptomic differences (Fig. 7). There were 6,779 differentially expressed genes (DEGs): 3,229 upregulated and 3,550 downregulated (Supplementary Material S5). The annotated volcano plot shows key DEGs (Fig. 8A): *PKIA*, *PLCB1*, and *EPDR1* were significantly downregulated in ESR1_12Z compared to NT_12Z and are involved in signal transduction and cell invasiveness (28–30). Unsurprisingly, *ESR1* was the most upregulated gene in the ESR1_12Z. Other notable upregulated genes include *TMPRSS15*, a protease that activates other proteases and cleaves proteins involved in cell signaling (31), and EPHA3, which inhibits cell migration (32) (Fig. 7A). To further characterize the DEGs, the KEGG (Kyoto Encyclopedia of Genes and Genomes) (33) dot plot shows the top 20 enriched pathways ordered by adjusted p-value (p_adj) with smaller values at the base (Fig. 7B). Out of the 14 pathways that were significantly enriched, several involved cellular signaling and cell-substrate adhesion functions such as cytokine-cytokine receptor interaction (p_adj = 2.40E-04), focal adhesion (p_adj = 6.69E-03), cell adhesion molecules (p_adj = 0.012), and ECM-receptor interaction (p_adj = 0.018). All of the significantly enriched KEGG pathways are provided for this comparison (Supplementary Material S6).

**Figure 7.**
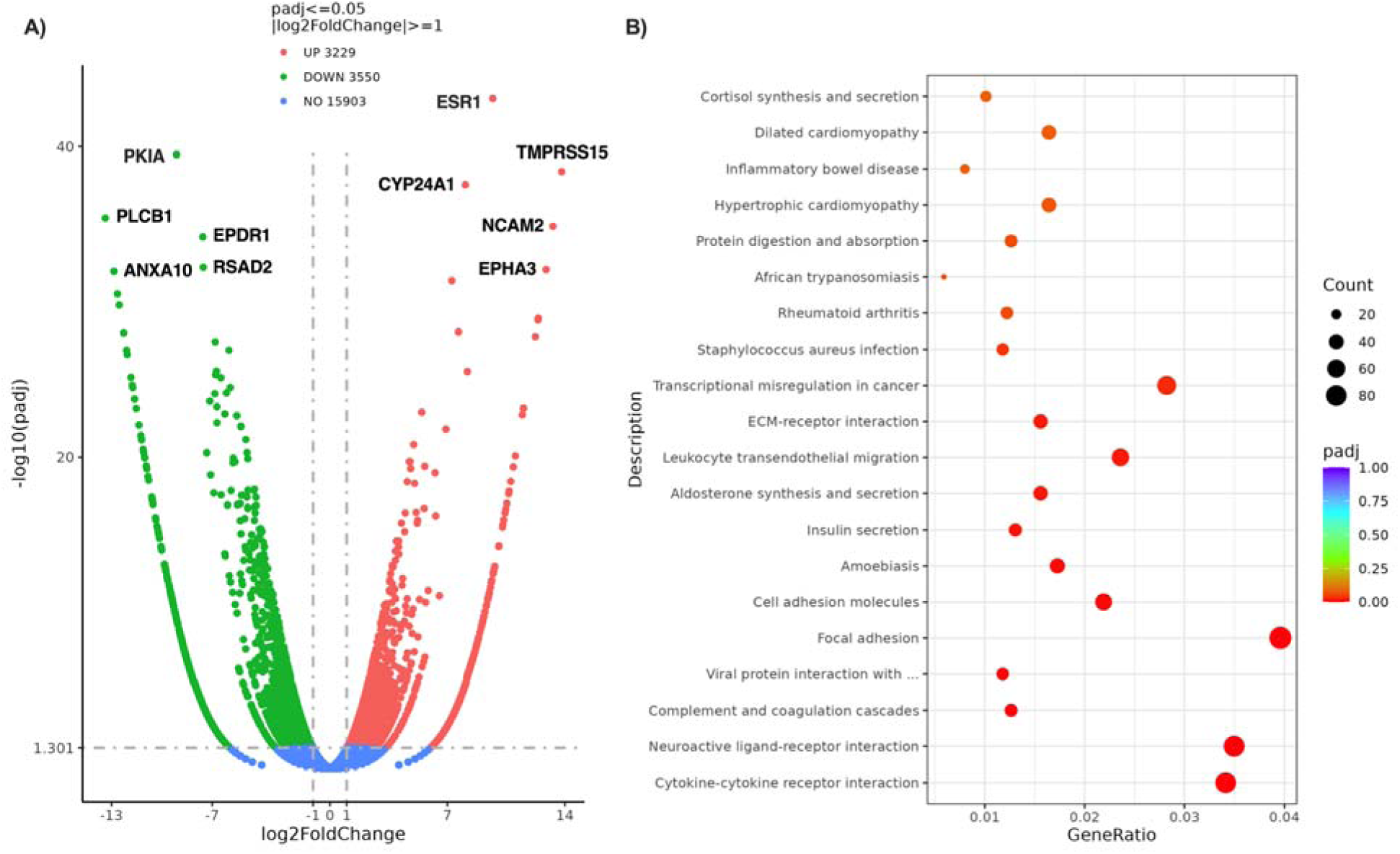
Differentially expressed genes and enriched pathways between 12Zs and *ESR1*-12Zs. **A)** The top 10 DEGs are annotated on the volcano plot, many of which are downregulated proteins involved in signal transduction and membrane trafficking. **B)** The top 20 enriched KEGG pathways are shown in the dot plot (ordered by p_adj, smaller values at the bottom), some of which function in cell signaling and cell-substrate adhesion. Differential gene expression analysis was carried out using EdgeR’s trimmed mean of M values with p_adj < 0.05 (24). KEGG (33) pathways with p_adj < 0.05 are considered significant. P-values were adjusted using the Benjamini-Hochberg method to control the error discovery rate (25).

## DISCUSSION

Estradiol (E2), a potent estrogen molecule, plays a fundamental role in regulating diverse physiological processes and contributes to the development of both benign and malignant conditions. Despite its broad impact, the precise mechanisms by which E2 modulates cellular activities in different tissue contexts remain incompletely understood. As researchers increasingly turn to *in vitro* models to study estrogen-dependent processes, it is critical to ensure that the cell lines used retain physiologically relevant responsiveness to E2. Without this validation, conclusions cannot drawn about estrogenic effects.

In the present study, we sought to characterize the sensitivity of the 12Z human endometriotic epithelial cell line to E2. While this cell line is frequently used in the study of estrogen-mediated conditions like endometriosis, our results challenge its assumed responsiveness to E2 in common *in vitro* models. We observed no significant changes in proliferation, collective migration, or single-cell random migration upon treatment with 10 nM E2 for 48 hours (Fig. 2). Together, these data suggest that the 12Z cell line lacks broad sensitivity to E2 in standard 2D *in vitro* systems, contrary to the sensitivity seen in primary endometriotic epithelial cells (13,17,34–36). A previous study demonstrated that E2 treatment of 12Zs did not enhance their production of pro-inflammatory cytokines (37). While this is a different experimental system, it reflects a similar theme of a lack of responsiveness to E2 in the 12Z cell line. Another study found that E2 treatment increased the invasion of heterogeneous spheroids composed of 12Z cells and endometriotic stromal cells within Matrigel containing peritoneal mesothelial cells (38); however, it is unclear whether the 12Zs, the stromal cells, or both were hormone responsive. In fact, the stromal cells might have been more sensitive to E2-induced invasiveness considering they were localized to the leading edges of the spheroids (38).

Interestingly, our results showed that the 12Zs did demonstrate a selective E2 responsiveness in physically confined migration systems (Fig. 3). This suggests that E2 may modulate cell behavior in a context-dependent manner, particularly in environments that mimic mechanical constraints present *in vivo*. It is possible that mechanical cues or confinement activate signaling pathways or receptor conformations that enable a cellular response to E2 that is otherwise dormant in 2D systems. This phenomenon underscores the importance of considering physical context when evaluating hormone responsiveness *in vitro*. Moreover, others have shown a differential expression of estrogen-related genes in 3D compared to 2D (22). We observed that E2 treatment modified the gene expression profile in 12Z cells, with a majority of the top 10 DEGs being pseudogenes or non-coding RNAs (Fig. 6). While their exact functions remain unclear, these molecules are increasingly recognized as regulators of gene expression through competing endogenous RNA activity, epigenetic modulation, or chromatin remodeling (26,39). It is possible that these upregulated molecules play a role in the expression of cytoskeletal regulators, mechanotransducers, cell volume regulators, or nuclear envelope genes. Physical confinement itself may activate E2 sensitivity in the 12Z cells, another question worthy of future investigation. This differential gene expression analysis differs from a previous study that reported an upregulation of some genes related to cell proliferation, apoptosis, motility, and signaling upon E2 treatment of the 12Zs (40). These contrasting results could be due to the use of different methods (proteomic analysis with mass spectrometry versus transcriptomic analysis with RNA sequencing in this study), different treatment incubation times (24 vs 48 hr), or different passage numbers.

Previously under the hypothesis that cellular sensitivity to E2 depends on *ESR1* expression, it seemed counterintuitive that although the overexpression of *ESR1* substantially increased ER⍰ protein levels, it did not translate to increased functional responsiveness to E2 in proliferation or migration assays (Fig. 4). This implies that E2 insensitivity in the 12Zs is not solely due to low or absent ER⍰ levels but may reflect downstream desensitization of estrogen signaling pathways or epigenetic modifications that uncouple ERs from functional outcomes. RNA sequencing of *ESR1*-12Zs revealed a downregulation of several key genes involved in signal transduction and membrane trafficking (e.g., *PKIA*, *PLCB1*, *EPDR1*, *ANXA10*) (Fig. 7), which may contribute to this lack of E2 sensitivity despite *ESR1* overexpression. Enriched KEGG pathways involving cytokine interactions, focal adhesions, cell adhesion molecules, and ECM-receptor signaling further support this hypothesis and suggest an altered regulatory network in *ESR1*-12Zs. Given the discrepancy between ER⍰ presence and the lack of functional response to E2, future studies should investigate 1) the balance between ER⍰ and ERβ in determining cellular E2 sensitivity and 2) the balance between classical (genomic) and non-classical (non-genomic) ER signaling pathways in the 12Zs. Classical ER signaling involves the direct binding of ligand-bound ERs to DNA at estrogen response elements (EREs), while non-classical signaling typically occurs through membrane-bound ERs that activate kinase cascades and secondary messengers (1). It is possible that the 12Z cells rely more heavily on non-genomic ER pathways that are context-dependent or that these signaling routes are impaired altogether. Dissecting the roles and integrity of these signaling pathways could provide insight into why some cellular behaviors, such as confined migration, are selectively responsive to E2 while others are not. Additionally, probing the involvement of G protein-coupled estrogen receptor or cytoplasmic ER pools could help clarify alternative routes of E2 influence.

Overall, these findings prompt a reevaluation of the 12Z cell line’s suitability as a model for estrogen-dependent processes. While it may still serve as a tool in endometriosis research, its limited E2 responsiveness in most *in vitro* contexts raises concerns about its utility for studying canonical estrogen signaling pathways. Future work could benefit from comparing the 12Z cell line to primary cell models with better-maintained hormonal sensitivity. Additionally, combining 12Zs with endometriotic stromal cells in co-culture, or subjecting cells to mechanical cues that mimic the *in vivo* microenvironment, may reveal hidden aspects of hormone signaling not captured in standard assays.

Our study has a few limitations. One limitation is that we only used endometriotic epithelial cells; the disease that the 12Z cell line models, endometriosis, is characterized by the presence of both glandular epithelium and stromal cells. It is unclear whether the presence of stromal cells contributes to the sensitivity of epithelial endometriotic cells to E2. Currently, there is no well-established endometriotic stromal cell line. However, future studies should consider testing E2 sensitivity in a co-culture of 12Zs and primary endometriotic stromal cells compared to each cell type individually. Another limitation of our study is that we obtained our 12Z cell line at passage 55. It is unknown how long the 12Z cell line can be passaged before it starts losing its original phenotype, but some changes have likely occurred: we detected nearly absent levels of ER⍰ in the 12Zs (Fig. 4A,B), contrary to earlier reports of ER⍰ expression in the 12Zs (6,7). We did not have access to earlier passages of the 12Zs, but it would be beneficial for others who do to study the effects of passaging on the retention of key traits of this particular cell line.

In conclusion, while 12Z cells are a widely used *in vitro* model of endometriosis, our findings indicate a restricted and context-specific responsiveness to E2. This selective behavior, especially under physical confinement, opens interesting avenues for further exploration into how mechanical and hormonal signaling intersect. However, researchers should exercise caution when using this cell line to model estrogen-dependent processes, particularly in 2D culture systems, and consider verifying hormone sensitivity in their specific experimental context.

## Supporting information

RNA sequencing methods

RNA sequencing Table 1

RNA sequencing Table 2

RNA sequencing Table 3

## AUTHOR CONTRIBUTIONS

KMS was the principal investigator. SB contributed to experimental design, data collection, data analysis, data interpretation, and manuscript preparation. EC contributed to data collection and data analysis. IMS contributed to data collection. All authors critically reviewed the manuscript.

## ACKNOWLEDGEMENTS

Biorender.com was used to create the schematic in Fig. 3C.

