## Supplementary material for "Selective estradiol sensitivity in 12Z human endometriotic epithelial cell line": RNA sequencing methods

### 1. Experimental Procedure

#### 1.1 Sample quality control

Please refer to QC report for methods of sample quality control.

#### 1.2 Library preparation for Transcriptome sequencing

##### Non strand specific library

Messenger RNA was purified from total RNA using poly-T oligo-attached magnetic beads. After fragmentation, the first strand cDNA was synthesized using random hexamer primers followed by the second strand cDNA synthesis. The library was ready after end repair, A-tailing, adapter ligation, size selection, amplification, and purification. The library was checked with Qubit and real-time PCR for quantification and bioanalyzer for size distribution detection.

##### Strand specific library

Messenger RNA was purified from total RNA using poly-T oligo-attached magnetic beads. After fragmentation, the first strand cDNA was synthesized using random hexamer primers. Then the second strand cDNA was synthesized using dUTP, instead of dTTP. The directional library was ready after end repair, A-tailing, adapter ligation, size selection, USER enzyme digestion, amplification, and purification. The library was checked with Qubit and real-time PCR for quantification and bioanalyzer for size distribution detection.

#### 1.3 Clustering and sequencing

After library quality control, different libraries were pooled based on the effective concentration and targeted data amount, then subjected to Illumina sequencing. The basic principle of sequencing is "Sequencing by Synthesis", where fluorescently labeled dNTPs, DNA polymerase, and adapter primers are added to the sequencing flow cell for amplification. As each sequencing cluster extends its complementary strand, the addition of each fluorescently labeled dNTP releases a corresponding fluorescence signal. The sequencer captures these fluorescence signals and converts them into sequencing peaks through computer software, thereby obtaining the sequence information of the target fragment.

#### 2. Bioinformatics Analysis Pipeline

##### 2.1 Data quality control

Raw data (raw reads) of fastq format were firstly processed through fastp software. In this step, clean data (clean reads) were obtained by removing reads containing adapter, reads containing ploy-N and low quality reads from raw data. At the same time, Q20, Q30 and GC content the clean data were calculated. All the downstream analyses were based on the clean data with high quality.

##### 2.2 Reads mapping to the reference genome

Reference genome and gene model annotation files were downloaded from genome website. Use HISAT2 (2.2.1) to build the index of the reference genome, and use HISAT2 to align paired-end clean reads to the reference genome. HISAT2 can use the gene model annotation file to create splice-aware alignments, providing better alignment accuracy compared to other non-splice alignment tools.

**2.3 Quantification of gene expression level**

featureCounts (2.0.6) was used to count the reads numbers mapped to each gene. And then FPKM of each gene was calculated based on the length of the gene and reads count mapped to this gene. FPKM, expected number of Fragments Per Kilobase of transcript sequence per Millions base pairs sequenced, considers the effect of sequencing depth and gene length for the reads count at the same time, and is currently the most commonly used method for estimating gene expression levels.

**2.4 Differential expression analysis**

For DESeq2 with biological replicates: Differential expression analysis for two conditions/groups was performed using the DESeq2 R package (1.42.0). DESeq2 provides statistical programs for determining differential expression in digital gene expression data using models based on negative binomial distribution. The resulting P-value is adjusted using the Benjamini and Hochberg’s methods to control the error discovery rate. The threshold of significant differential expression: padj <= 0.05 & |log2(foldchange)| >= 1.

For edgeR without biological replicates: Prior to differential gene expression analysis, for each sequencing library, read counts were adjusted using the edgeR R package (4.0.16) by scaling normalization factors to eliminate differences in sequencing depth between samples, followed by differential expression analysis. The resulting P value is adjusted using the Benjamini and Hochberg’ s methods to control the error discovery rate. The threshold of significant differential expression: padj <= 0.05 & |log2(foldchange)| >= 1.

**2.5 GO and KEGG enrichment analysis of differentially expressed genes**

Gene Ontology (GO) enrichment analysis of differentially expressed genes was implemented by the clusterProfiler (4.8.1), in which gene length bias was corrected. GO terms with corrected P-value less than 0.05 were considered significantly enriched by differential expressed genes. KEGG is a database resource for understanding high-level functions and utilities of the biological system, such as the cell, the organism and the ecosystem, from molecular-level information, especially large-scale molecular datasets generated by genome sequencing and other high-through put experimental technologies (http://www.genome.jp/kegg/). We used clusterProfiler R package to test the statistical enrichment of differential expression genes in KEGG pathways.

**2.6 Gene Set Enrichment Analysis**

Gene Set Enrichment Analysis (GSEA) is a computational approach to determine if a predefined Gene Set can show a significant consistent difference between two biological states. The genes were ranked according to the degree of differential expression in the two samples, and then the predefined Gene Set were tested to see if they were enriched at the top or bottom of the list. Gene set enrichment analysis can include subtle expression changes. We use the local version of the GSEA analysis tool http://www.broadinstitute.org/gsea/index.jsp, GO, KEGG data set were used for GSEA independently.

**2.7 PPI analysis of differentially expressed genes**

PPI analysis of differentially expressed genes was based on the STRING database, which contains known and predicted protein-protein interactions. For species present in the database, we construct the network by extracting the target gene list from the database. Otherwise, we use diamond (version 0.9.13) to align the target gene sequences with selected reference protein sequences, and then establish the network based on the known interactions of the selected reference species.

**2.8 AS analysis**

Alternative Splicing is an important mechanism for regulating gene expression and protein variability. The rMATS (4.3.0) software was used to analyze alternative splicing (AS) events, which mainly include five types of variable splicing events: SE (skipped exon), RI (retained intron), MXE (mutually exclusive exons), A5SS (5' alternative splice site), and A3SS (3' alternative splice site).

##### 2.9 SNP analysis

The GATK (v4.1.1.0) software was used to perform SNP calling.

##### 2.10 Fusion Analysis

Fusion gene refers to the chimeric gene formed by the fusion of all or part of the sequences of two genes, which is generally caused by chromosome translocation, deletion and other reasons. The STAR_fusion (version 1.12) software was utilized for the detection of fusion genes. SATR_fusion is a software package uses fusion output results of STAR alignment to detect fusion transcripts, including SATR alignment, SATRfusion.predict, SATRfusion.filter was used to correct the predicted results of SATR_fusion to ensure the accuracy of the results.
