## Supplementary Figure S2 for "Selective estradiol sensitivity in 12Z human endometriotic epithelial cell line"

**SUPPLEMENTARY MATERIAL S2**

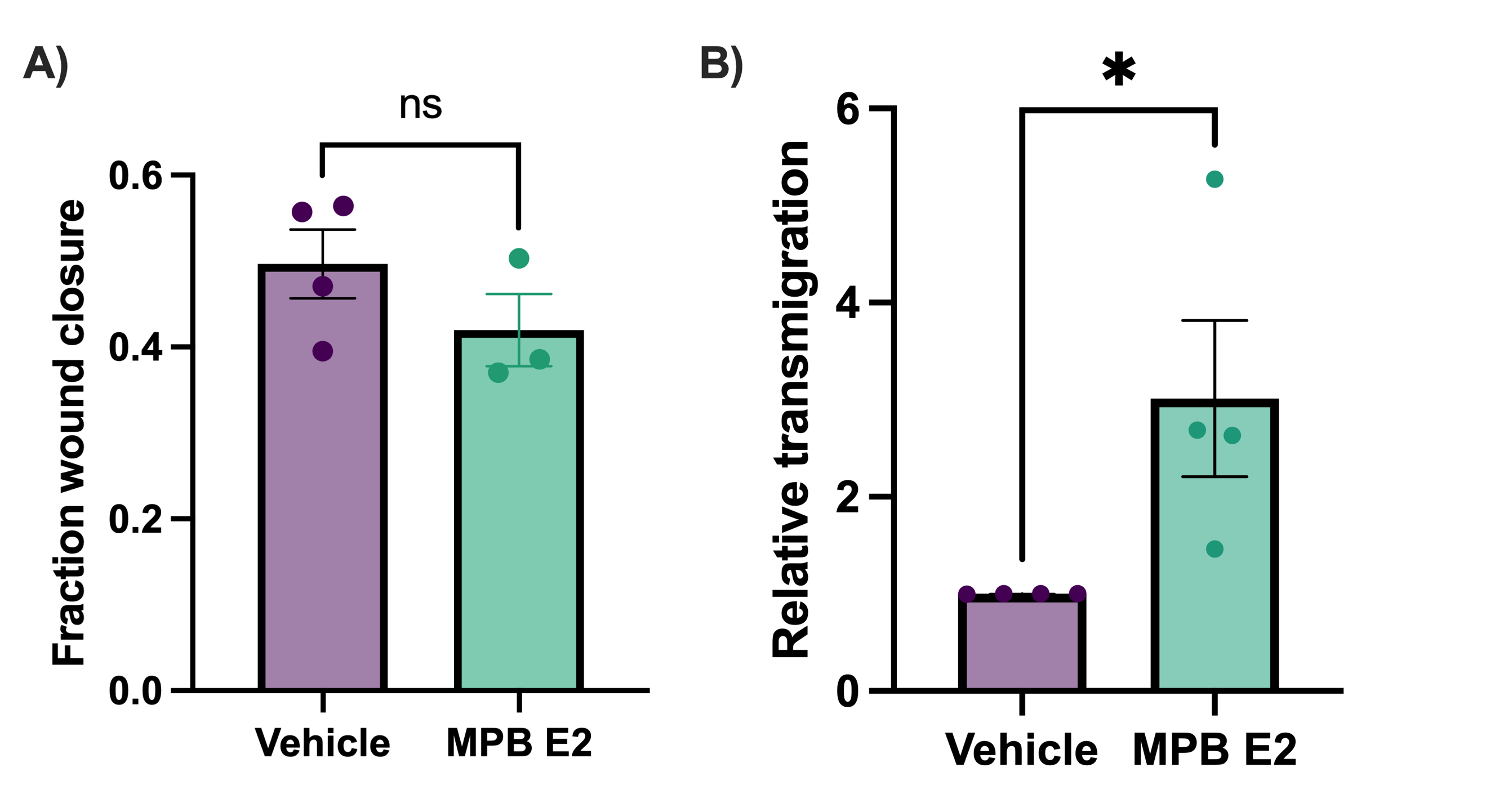
We found that the 12Z cell line demonstrated a lack of responsiveness to E2 treatment in proliferation and migration assays (Fig. 2) which prompted us to question the reliability of our source of E2. We obtained a new vial of 17β-estradiol (MP Biomedicals), denoted MPB E2. We treated 12Zs with 10 nM MPB E2 or the vehicle control (0.001% EtOH) for 48 hr and tested their migratory response. MPB E2 had no effect on wound closure (Fig. S2A) but did significantly increase transmigration in a transwell assay (Fig. S2B). These trends matched our results using the original source of E2 (Sigma Aldrich, cat. #E8875-1G), as seen in Fig. 2B and Fig. 5A. Therefore, we found no reason to question the integrity of the E2 from Sigma Aldrich, whose Certificate of Analysis reports a purity greater than 98% by high-performance liquid chromatography.

**Figure S2. Functional assessment of alternative source of E2 on 12Zs.** E2 purchased from MP Biomedicals (MPB E2) was applied to 12Zs and their migratory response was determined. **A)** Fraction of wound closed is reported and **B)** transwell migration is reported relative to the vehicle control. Data points represent biological replicates, error bars represent standard error of the mean. *P ≤ 0.05. MPB E2; MP Biomedicals estradiol.
