## Supplementary Figure S3 for "Selective estradiol sensitivity in 12Z human endometriotic epithelial cell line"

**SUPPLEMENTARY MATERIAL S3**

*ESR1*-12Zs acquired from Dr. Fazleabas’s lab (Michigan State University) were tested against 12Zs to verify the upregulation of ER⍺. We also tested the effect of E2 on ER⍺ expression in both cell lines. Western blots were performed in triplicate (Fig. S3A). E2 had no effect on ER⍺ expression in 12Zs (Fig. S3B,A) or *ESR1*-12Zs (Fig. S3B,B).

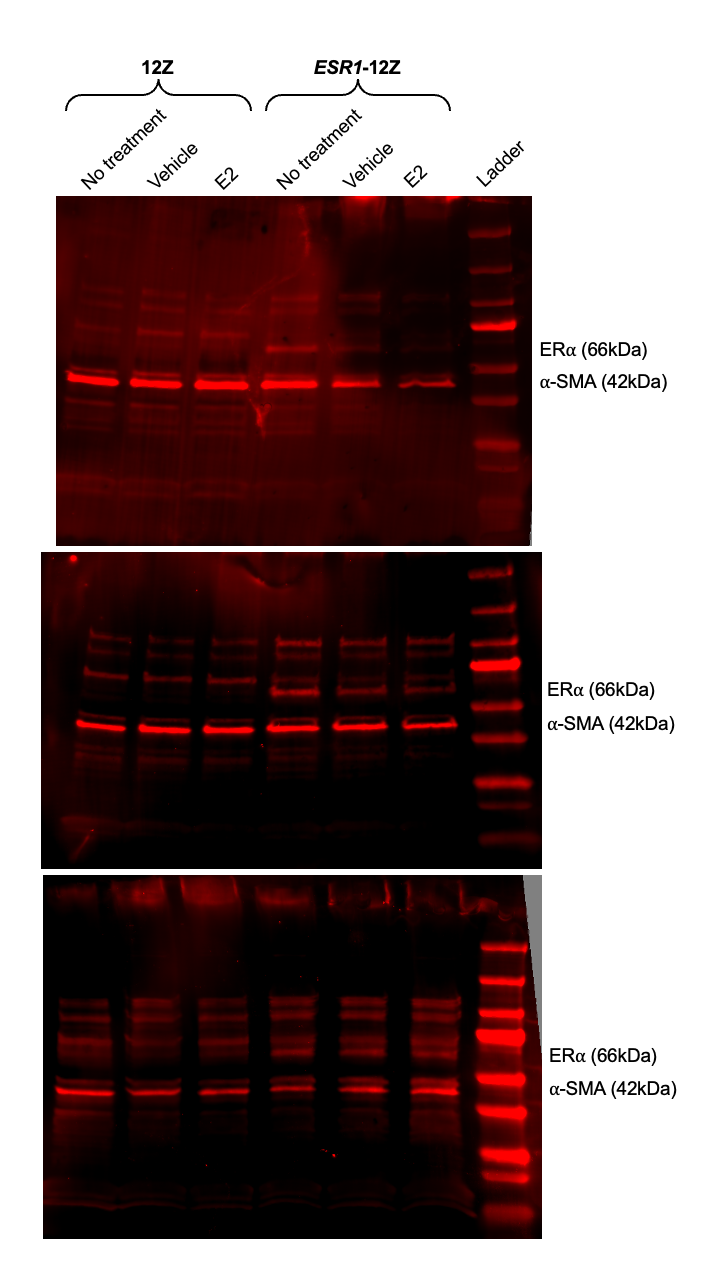
**Figure S3A. Western blots measuring upregulation of ER⍺ in *ESR1*-12Zs.** Western blots were performed using ⍺-SMA (42kDa) as the reference proztein and the SeeBlue Plus2 prestained protein ladder. Quantification of ER⍺ expression in *ESR1*-12Zs and 12Zs (no treatments) is shown in the main article in Fig. 4B. Blots are from three independent trials. E2; estradiol, ER⍺; estrogen receptor-⍺, ⍺-SMA; ⍺-smooth muscle actin.

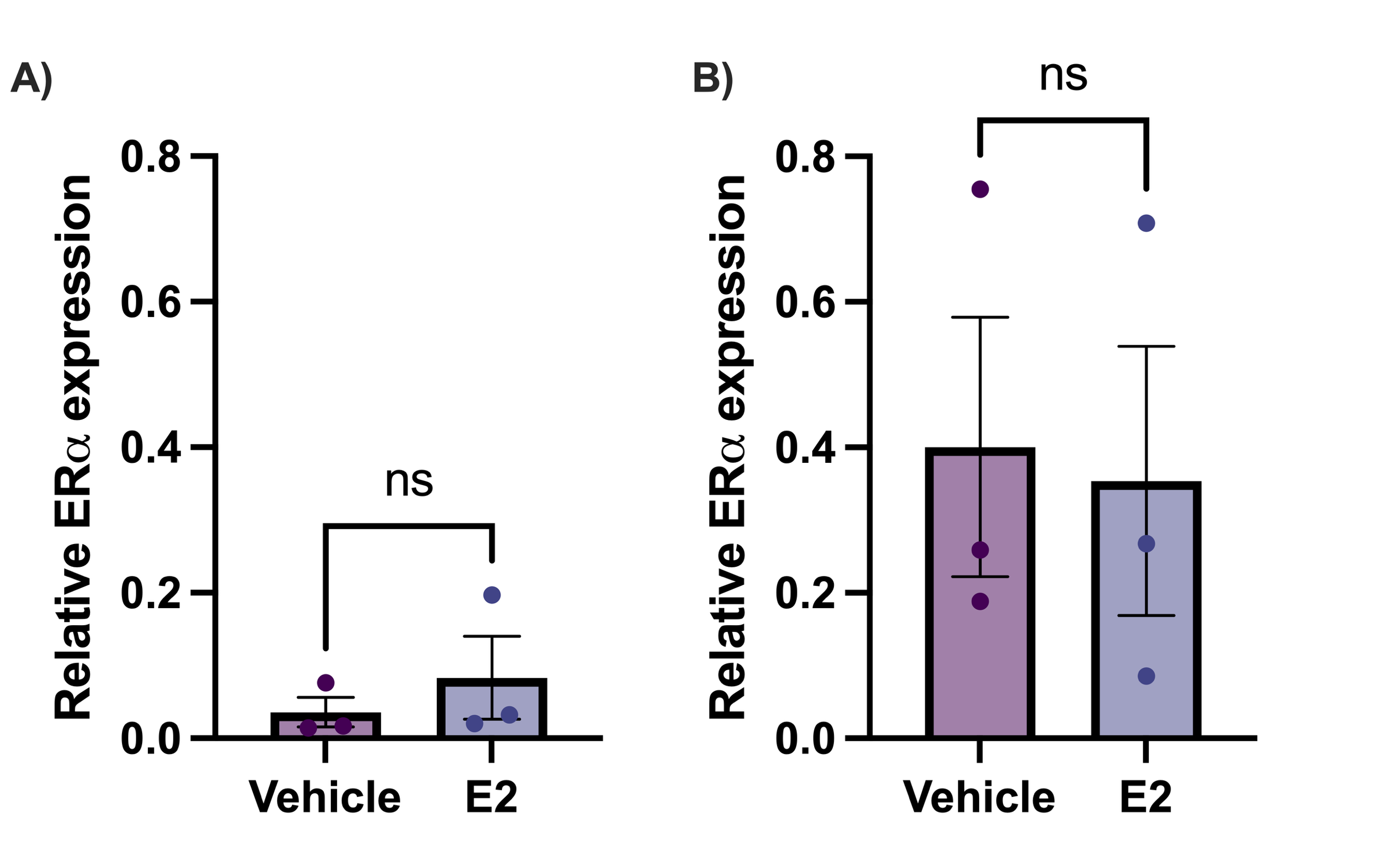
**Figure S3B. Effect of E2 on ER⍺ expression in 12Zs and *ESR1*-12Zs.** Cells were treated with vehicle (0.001% EtOH) and 10nM E2 for 48 hr prior to lysis. ER⍺ expression was quantified relative to ⍺-SMA expression in **A)** 12Zs and **B)** *ESR1*-12Zs. Data points represent biological replicates, error bars represent standard error of the mean. E2; estradiol.
